# Ablation of Mitochondrial RCC1-L Induces Nigral Dopaminergic Neurodegeneration and Parkinsonian-like Motor Symptoms

**DOI:** 10.1101/2023.12.01.567409

**Authors:** Kaylin J. Ellioff, Susan M.K. Osting, Alyssa Lentine, Ashley D. Welper, Corinna Burger, Daniel S. Greenspan

## Abstract

Mitochondrial dysfunction has been linked to both idiopathic and familial forms of Parkinson’s disease (PD). We have previously identified RCC1-like (RCC1L) as a protein of the inner mitochondrial membrane important to mitochondrial fusion. Herein, to test whether deficits in RCC1L mitochondrial function might be involved in PD pathology, we have selectively ablated the *Rcc1l* gene in the dopaminergic (DA) neurons of mice. A PD-like phenotype resulted that includes progressive movement abnormalities, paralleled by progressive degeneration of the nigrostriatal tract. Experimental and control groups were examined at 2, 3-4, and 5-6 months of age. Animals were tested in the open field task to quantify anxiety, exploratory drive, locomotion, and immobility; and in the cylinder test to quantify rearing behavior. Beginning at 3-4 months, both female and male *Rcc1l* knockout mice show rigid muscles and resting tremor, kyphosis and a growth deficit compared with heterozygous or wild type littermate controls. *Rcc1l* knockout mice begin showing locomotor impairments at 3-4 months, which progress until 5-6 months of age, at which age the *Rcc1l* knockout mice die. The progressive motor impairments were associated with progressive and significantly reduced tyrosine hydroxylase immunoreactivity in the substantia nigra pars compacta (SNc), and dramatic loss of nigral DA projections in the striatum. Dystrophic spherical mitochondria are apparent in the soma of SNc neurons in *Rcc1l* knockout mice as early as 1.5-2.5 months of age and become progressively more pronounced until 5-6 months. Together, the results reveal the RCC1L protein to be essential to *in vivo* mitochondrial function in DA neurons. Further characterization of this mouse model will determine whether it represents a new model for *in vivo* study of PD, and the putative role of the human *RCC1L* gene as a risk factor that might increase PD occurrence and severity in humans.

## INTRODUCTION

Parkinson’s disease (PD), a progressive neurodegenerative disorder, is the second most common neurodegenerative disorder after Alzheimer’s disease (Alzheimer’s, 2014; Dorsey et al., 2007), and affects ∼1-3% of all adults over age 65 (de Rijk et al., 2000; Dorsey et al., 2007). PD is characterized by dysfunctions of the somatomotor system that include bradykinesia, muscular rigidity, postural imbalance, gait dysfunction, and resting tremor (Gibb and Lees, 1988; Litvan et al., 2003). The pathological hallmark of the PD motor dysfunction phenotype is progressive degeneration of the nigrostriatal dopaminergic pathway, with progressive death of substantia nigra pars compacta (SNc) dopaminergic (DA) neurons and depletion of dopamine in the basal ganglia (Dauer and Przedborski, 2003; Mullin and Schapira, 2015). Mitochondrial functional deficits have for some time been associated with PD, as compounds that inhibit the respiratory chain can cause PD-like symptoms, and respiratory chain deficits have been identified in PD patients (Chen et al., 2019). In addition, although ∼90% of PD cases are sporadic, with no clearly identified single cause, ∼10% of cases have a clear family history of PD due to mutations in single genes that result in mitochondrial dysfunction and defects in lysosomal and autophagy pathways (Li et al., 2021; Nechushtai et al., 2023; Rakovic et al., 2011). Gene products associated with familial, monogenic PD include Parkin (a mitochondrially located E3 ubiquitin ligase with roles in mitochondrial quality control), PINK1 (a mitochondrially located kinase activator of Parkin), DJ-1 (localized to mitochondria under conditions of oxidative stress and protective against reactive oxygen species, ROS), LRRK2 (regulates activities of mitochondrially localized E3 ubiquitin ligases to control mitochondrial-ER tethering/bioenergetics), and VPS35 (involved in controlling mitochondrial fission)(Clark et al., 2006; Greene et al., 2003; Irrcher et al., 2010; Krebiehl et al., 2010; Park et al., 2006; Tang et al., 2015; Wang et al., 2012).

Mitochondria normally undergo dynamic cycles of inter-mitochondrial fusion and fission that form and remodel tubular networks, thereby optimizing intracellular mitochondrial distribution, respiratory function, and control of apoptosis and autophagy, as well as preventing genetic drift in mitochondrial DNA mutation distribution (Chan, 2006; Kasahara and Scorrano, 2014). Mitochondrial fusion is driven by outer membrane GTPases MFN1 and MFN2 and inner membrane GTPase OPA1 (Chan, 2006). Mfn2 and OPA1 mutations are causal in the neural degenerative diseases Charcot-Marie-Tooth neuropathy type 2A (neuropathy of long motor and sensory neurons) (Zuchner et al., 2004) and dominant optic atrophy (optic nerve degeneration, ataxia, deafness, and peripheral neuropathy) (Delettre et al., 2000), respectively, demonstrating the importance of mitochondrial fusion to neuronal function *in vivo*. A link to PD was demonstrated by studies in which DA neuron-specific knockdown of *Mfn2* produced mice with PD-like phenotypes that included severe striatal loss of DA nerve terminals, and DA neuron loss in the SNc (Lee et al., 2012; Pham et al., 2012). In fact, Parkin normally targets MFN2 for proteasomal degradation by isolating damaged mitochondria for degradation via mitophagy, as part of the mitochondrial quality control mechanism (Twig et al., 2008). Parkin also appears to play a role in increasing OPA1 expression, thus helping stabilize mitochondrial cristae (Gegg et al., 2010; Rakovic et al., 2011). VPS35 mutations may impair mitochondrial fusion by stabilizing mitochondrial E3 ubiquitin ligase 1 (MUL1), thereby increasing MFN2 degradation and thus shifting the balance of fission/fusion towards increased fission/fragmentation (Tang et al., 2015).

Our group previously identified WBSCR16 as a protein of the inner mitochondrial membrane with importance to mitochondrial fusion (Huang et al., 2017). Additionally, a genome-wide screen by others (Arroyo et al., 2016) identified WBSCR16, among other gene products, as being important to oxidative phosphorylation, and association of WBSCR16 with the 16S rRNA of the mitochondrial ribosome (mitoribosome) with suggested roles in intramitochondrial protein translation. After these reports, the mouse *Wbscr16* gene was formally renamed *Rcc1l* and, thus, the gene and gene product shall be referred to as *Rcc1l* and RCC1L, respectively through the remainder of this report. RCC1L has also been found to play a role in pseudouridylation, and stabilization of mitoribosomes (Antonicka et al., 2017; Kramer et al., 2023; Proust et al., 2021; Reyes et al., 2020). Relevant to human health, RCC1L has been identified as a biomarker for predicting the prognosis of adrenal cortical carcinoma (Jang et al., 2021).

We previously showed homozygosity for a spontaneous missense mutation of the *Rcc1l* gene to cause embryonic lethality, apparently due to deficits in early placentation. E*x vivo* studies revealed that neurons of mice heterozygous for this mutation have mitochondria with reduced membrane potential and increased susceptibility to fragmentation upon excitotoxic stress (Huang et al., 2017). Mitochondria are essential for energy provision in all eukaryotic cells, and optimal mitochondrial function is especially crucial to neurons, due to their high energy requirements. In particular, DA neurons depend on optimal mitochondrial function for the especially high energy requirements to clear ROS produced by dopamine metabolism (Durcan and Fon, 2015). These mitochondrial requirements of DA neurons doubtless underlie the association of deficits in mitochondrial function with PD (above). Here, we have utilized the sensitivity of DA neurons to mitochondrial dysfunction and the relationship of DA neuronal dysfunction to PD to gauge the extent to which RCC1L loss might affect mitochondrial function *in vivo*. Towards this end, we have employed DA neuron-specific *Rcc1l* knockout in mice and characterized a resulting PD-like phenotype that includes progressive movement abnormalities and degeneration of the nigrostriatal circuit. Results herein demonstrate an important *in vivo* role for RCC1L in mitochondrial function in DA neurons.

## RESULTS

### Generation of mice for conditional ablation of *Rcc1l* in DA neurons

We have previously shown that homozygosity for *Wbscr16*/*Rcc1l* mutations can be early embryonic lethal, with conceptus growth arrest at E8.5 (Huang et al., 2017). To determine the effects of *Rcc1l* ablation *in vivo*, post E8.5, we began study of mice with a targeted *Rcc1l* allele in which *Rcc1l* exon 2 is floxed and *Rcc1l* intron 1 contains a *lacZ* cassette (Fig. 1A). Transcription of this targeted allele from the endogenous *Rcc1l* promoter yields a transcript comprising *Rcc1l* exon 1 spliced to the lacZ sequences, disrupting *Rcc1* such that it is a null allele. Consistent with the latter, progeny homozygous for this putatively null allele were not obtained in any of numerous litters (data not shown). To obtain a lacZ-minus, fully functional *Rcc1l* allele, the FRT flanked lacZ cassette was excised via crosses of mice with the targeted *Rcc1l* allele to ROSA26::FLPe mice, in which FLPe, is broadly expressed in tissues (including germ cells) (Farley et al., 2000)(Fig. 1A). The resulting functional *Rcc1lRcc1l*^*fl*/*fl*^ allele differs from wild type in having a floxed exon 2, allowing conditional *Rcc1l* inactivation by Cre recombinase.

**Figure 1.**
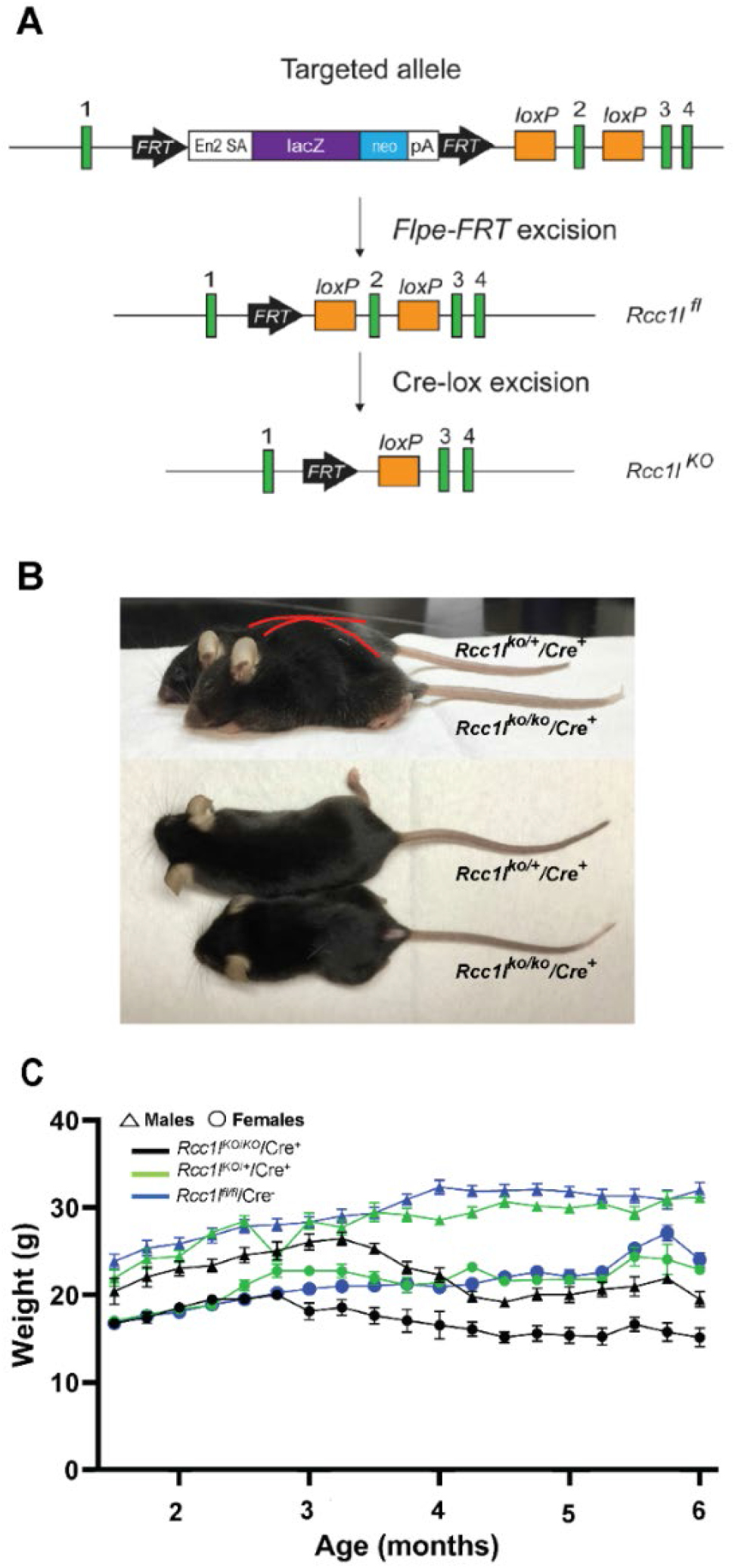
DA-neuron-specific knockdown of *Rcc1l* results in growth deficits and kyphosis in mice. (A) Schematic shows placement of loxP and FRT sites and lacZ cassette in targeted *Rcc1l* allele and shows effects of Flpe- and Cre-induced excisions. Exons, green boxes. *Rccl1* exon 2 is flanked by *loxP* sites, such that excision yields loss of 130 bp of coding sequence and a frameshift, so that only 76% of RLD repeat 1 is transcribed, fused to 37 amino acids in a new reading frame, prior to a premature stop. Other RLD repeats are lost, and mutant RNA may not be stable due to nonsense-mediated decay. In the targeted allele, En2 SA is an acceptor site for splicing of Rcc1l exon 1 to lacZ sequences, pA is the polyadenylation site for the lacZ cassette. *Rcc1l*^*fl*^, allele with excised lacZ cassette and floxed exon 2; *Rcc1l*^*KO*^, allele in which both lacZ cassette and exon2 have been excised. (B) Representative images of 21-week-old female *Rcc1l*^*KO*/*KO*^/*Cre*^+^ mice and heterozygous *Rcc1l*^*KO*/*+*^/*Cre*^+^controls. Red outlining highlights the differing dorsal contours of *Rcc1l*^*KO*/*KO*^/*Cre*^+^ and *Rcc1l*^*KO*/*+*^/*Cre*^+^mice and kyphosis of the former. (C) Graph of weekly weight measurements of male and female *Rcc1l*^*KO*/*KO*^/*Cre*^+^ (KO), *Rcc1l*^*KO*/*+*^/*Cre*^+^ (Het), and *Rcc1l*^*fl*/*fl*^/*Cre*^-^ (*Cre*-) mice, in which each point is the average weight ± SEM. Both male and female *Rcc1l*^*KO*/*KO*^/*Cre*^+^ mice are significantly smaller than control animals by 3.5 months of age.

To directly determine whether RCC1L is important to DA neuronal function, mice heterozygous for the *Rcc1l*^*fl*/*fl*^ locus were crossed with mice with the *Slc6a3*-*Cre* loci, in which the Cre transgene has been knocked into the endogenous *Slc6a3* gene, which encodes the dopamine transporter (DAT), and in which Cre expression is driven by the endogenous *Slc6a3* cis-acting elements, such that Cre is expressed only in A8—A10 DA-neurons (Backman et al., 2006). Subsequently, *Rcc1l*^*fl*/*fl*^ heterozygotes with the *Slc6a3*-*Cre* transgene were crossed with *Rcc1l*^*fl*/*fl*^ heterozygotes lacking the *Slc6a3*-*Cre* transgene (Supplemental Fig.1).

### Loss of RCC1L produces an age-dependent movement disorder

Resulting *Rcc1l*^*KO*/*KO*^/Cre^+^ progeny, homozygous for exon 2-excised/null *Rcc1l* alleles, exhibited kyphosis and a growth deficit compared with control progeny either heterozygous for both the null allele and *Slc6a3*-*Cre* transgene (*Rcc1l*^*KO*/*+*^/Cre^+^) or homozygous for the floxed allele and Cre-negative (*Rcc1l*^*fl*/*fl*^/Cre^-^) (Fig. 1B). *Rcc1l*^*KO*/*KO*^/Cre^+^ mice die by ∼6 months of age. By 3-months of age, both female and male *Rcc1l*^*KO*/*KO*^/*Cre*^+^ mice showed significant growth deficit compared to *Rcc1l*^*fl*/*fl*^/*Cre*^-^ and *Rcc1l*^*KO*/*+*^/*Cre*^+^ (Figs. 1B and C). When looking at the mixed effects, genotype by time interaction was found in *Rcc1l*^*KO*/*KO*^/*Cre*^+^ relative to the *Rcc1l*^*fl*/*fl*^/*Cre*^-^ and *Rcc1l*^*KO*/*+*^/*Cre*^+^ controls (males: F _(36, 479)_ = 12.09, and females: F _(36, 544)_ = 7.371, p<0.0001). There was a slight sex difference in the growth deficit, as *Rcc1l*^*KO*/*KO*^/*Cre*^+^ female mice were already significantly smaller than controls by 13 weeks of age (*Rcc1l*^*KO*/*KO*^/*Cre*^+^ *vs. Rcc1l*^*fl*/*fl*^/*Cre*^-^ one way ANOVA main effect of treatment, F _(2, 33)_ = 11.83, p=0.0001; *Rcc1l*^*KO*/*KO*^/Cre^+^ *vs. Rcc1l*^*KO*/*+*^/*Cre*^+^, p=0.004) (Fig. 1C). Males started to show stunted growth at 15 weeks of age. In addition, at 3-4 months of age both female and male *Rcc1l*^*KO*/*KO*^/Cre^+^ mice started to show noticeable bradykinesia, rigid muscles, impaired posture, and tremor when at rest (not shown), which were not observed in *Rcc1l*^*KO*/*+*^/*Cre*^+^ or *Rcc1l*^*fl*/*fl*^/*Cre*^-^ control sibling mice.

To quantify the observed movement impairment of *Rcc1l*^*KO*/*KO*^/Cre^+^ mice they, together with *Rcc1l*^*KO*/*+*^/*Cre*^+^ and *Rcc1l*^*fl*/*fl*^/*Cre*^-^ controls, were subjected to open field testing of spontaneous locomotion at three different times in their lifespan (1.5-2.5 months of age, 3-4 months of age, and 5-6 months of age) (Fig. 2). *Rcc1l*^*KO*/*KO*^/*Cre*^+^ had a pronounced age-dependent reduction in longitudinal locomotion/increased immobility that was not seen in either of the controls (Fig.2A-C). At 3-4 months of age there was a statistically significant difference in the path travelled by *Rcc1l*^*KO*/*KO*^/*Cre*^+^ mice relative to *Rcc1l*^*KO*/*+*^/*Cre*^+^ and *Rcc1l*^*fl*/*fl*^/*Cre*^-^ controls (*Rcc1l*^*KO*/*KO*^/*Cre*^+^ vs. *Rcc1l*^*fl*/*fl*^/*Cre*^-^ one way ANOVA main effect of treatment, F_(2, 18)_ =25.44, p=0.0001; *Rcc1l*^*KO*/*KO*^/*Cre*^+^ vs. *Rcc1l*^*KO*/*+*^/*Cre*^+^, p<0.0001). At 5-6 months, *Rcc1l*^*KO*/*KO*^/*Cre*^+^ mice were scarcely moving (Fig. 2B and C) (*Rcc1l*^*KO*/*KO*^/*Cre*^+^ vs. *Rcc1l*^*fl*/*fl*^/*Cre*^-^ main effect of treatment, F_(2, 11)_ =37.23, p=0.0002; *Rcc1l*^*KO*/*KO*^/*Cre*^+^ vs. *Rcc1l*^*KO*/*+*^/*Cre*^+^, p<0.0001). We also measured increased immobility, which was apparent in *Rcc1l*^*KO*/*KO*^/*Cre*^+^ mice. Here, we saw increased immobility beginning at 3-4 months (*Rcc1l*^*KO*/*KO*^/*Cre*^+^ vs. *Rcc1l*^*fl*/*fl*^/*Cre*^-^ one way ANOVA main effect of treatment, F_(2, 11)_ =7.262, p=0.001; *Rcc1l*^*KO*/*KO*^/*Cre*^+^ vs. *Rcc1l*^*KO*/*+*^/*Cre*^+^, p=0.04) and at 5-6 months (*Rcc1l*^*KO*/*KO*^/*Cre*^+^ vs. *Rcc1l*^*fl*/*fl*^/*Cre*^-^ one way ANOVA main effect of treatment, F_(2, 9)_ =7.09, p=0.03; *Rcc1l*^*KO*/*KO*^/*Cre*^+^ vs. *Rcc1l*^*KO*/*+*^/*Cre*^+^, p=0.02). A cylinder test of forelimb akinesia showed *Rcc1l*^*KO*/*KO*^/*Cre*^+^ mice to have a profound deficiency in rearing behavior that progressed with age (Fig. 2D). At 3-4 months, *Rcc1l*^*KO*/*KO*^/*Cre*^+^ mice rear significantly less compared with control groups (*Rcc1l*^*KO*/*KO*^/*Cre*^+^ vs. *Rcc1l*^*fl*/*fl*^/*Cre*^-^ one way ANOVA main effect of treatment, F_(2, 15)_ =20.41, p=0.002; *Rcc1l*^*KO*/*KO*^/*Cre*^+^ vs. *Rcc1l*^*KO*/*+*^/*Cre*^+^, p<0.0001). At 5-6 months this difference becomes more pronounced (*Rcc1l*^*KO*/*KO*^/*Cre*^+^ vs. *Rcc1l*^*fl*/*fl*^/*Cre*^-^ one way ANOVA main effect of treatment, F_(2, 12)_ =73.54, p<0.0001; *Rcc1l*^*KO*/*KO*^/*Cre*^+^ vs. *Rcc1l*^*KO*/*+*^/*Cre*^+^, p<0.0001). A significant difference in the number of rears between *Rcc1l*^*KO*/*+*^/*Cre*^+^ and *Rcc1l*^*fl*/*fl*^/*Cre*^-^ control groups was observed at 5-6 months of age (*Rcc1l*^*KO*/*+*^/*Cre*^+^ vs. *Rcc1l*^*fl*/*fl*^/*Cre*^-^ one way ANOVA main effect of treatment, F_(2, 12)_=73.54, p=0.003).

**Figure 2.**
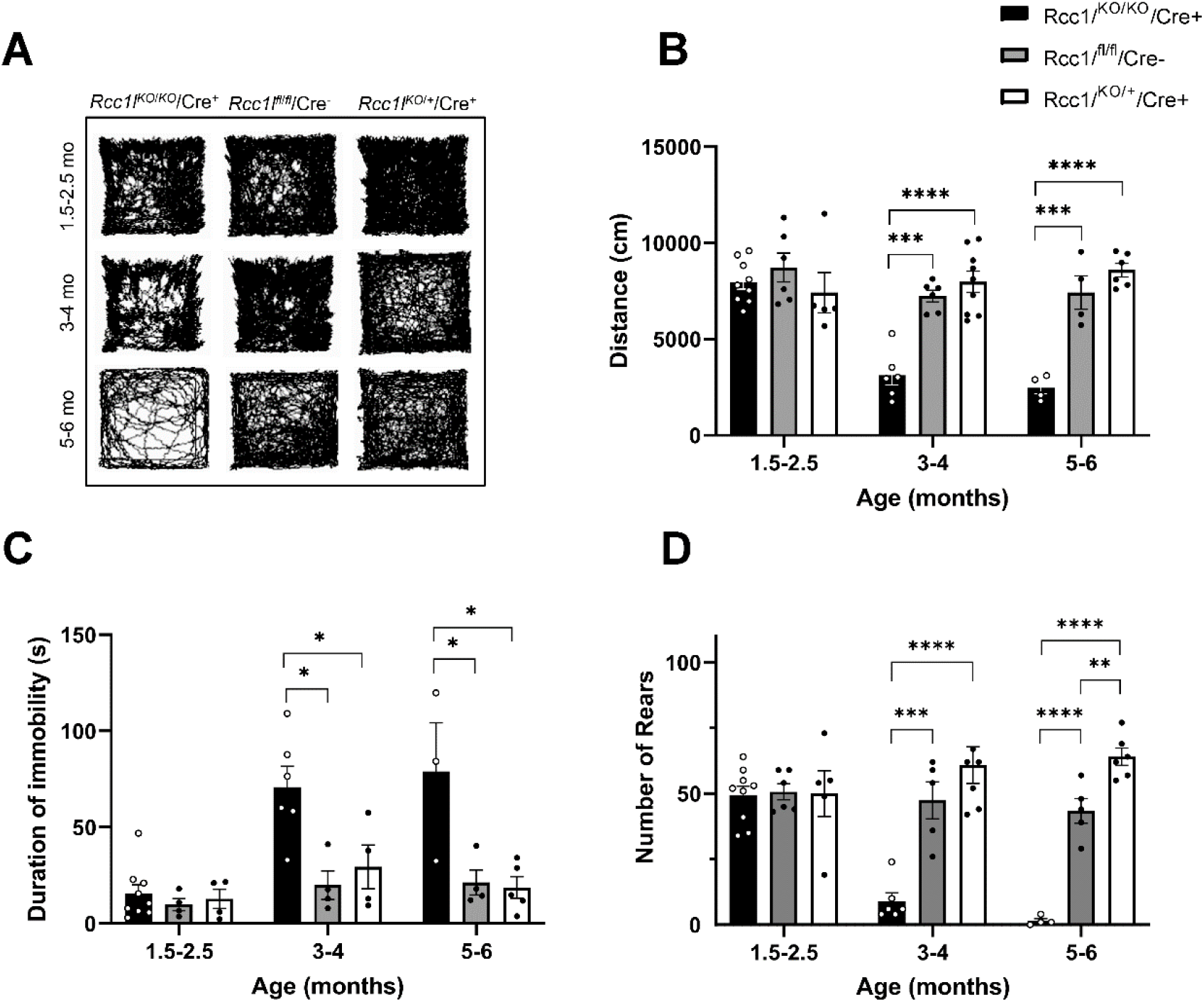
*Rcc1l*^*KO*/*KO*^/Cre^+^ mice experience age-dependent reduction in motor function. (A) Representative traces of open field analysis of movement that occurred over a 15 min period. (B) Quantification of distance traveled in the open field analysis. (C) Duration of immobility. (D) Quantification of rearing in the cylinder assay. *, p<0.05; **, p<0.01; ***, p<0.001; ****, p<0.0001

### *Rcc1l*-null mutants display an age-dependent degeneration of nigrostriatal dopaminergic neurons

DA neuron-specific knockdown of Mfn2 transgenic mice led to severe DA neuron loss in the substantia nigra pars compacta (SNc), as well as loss of DA nerve terminals in the striatum (Lee et al., 2012; Pham et al., 2012). Thus, we assayed for possible DA loss in the nigrostriatal circuit in *Rcc1l*-null mice by examining the expression of tyrosine hydroxylase (TH), one of the limiting enzymes in the production of DA in the SNc. To determine the anatomical distribution of TH, protein expression was visualized using immunohistochemistry in the SNc and striatum of the three experimental groups. Beginning at 3-4 months of age *Rcc1l*^*KO*/*KO*^/*Cre*^+^ display significant decrease in TH immunoreactivity in the SNc compared to *Rcc1l*^*fl*/*fl*^/*Cre*^-^ and *Rcc1l*^*fl*/*fl*^/*Cre*^-^ controls (Fig 3 A, and supplemental figures 2-4 for rostro caudal cross-sectional series). This decrease becomes more apparent at 5-6 months of age (Fig. 3A).

**Figure 3.**
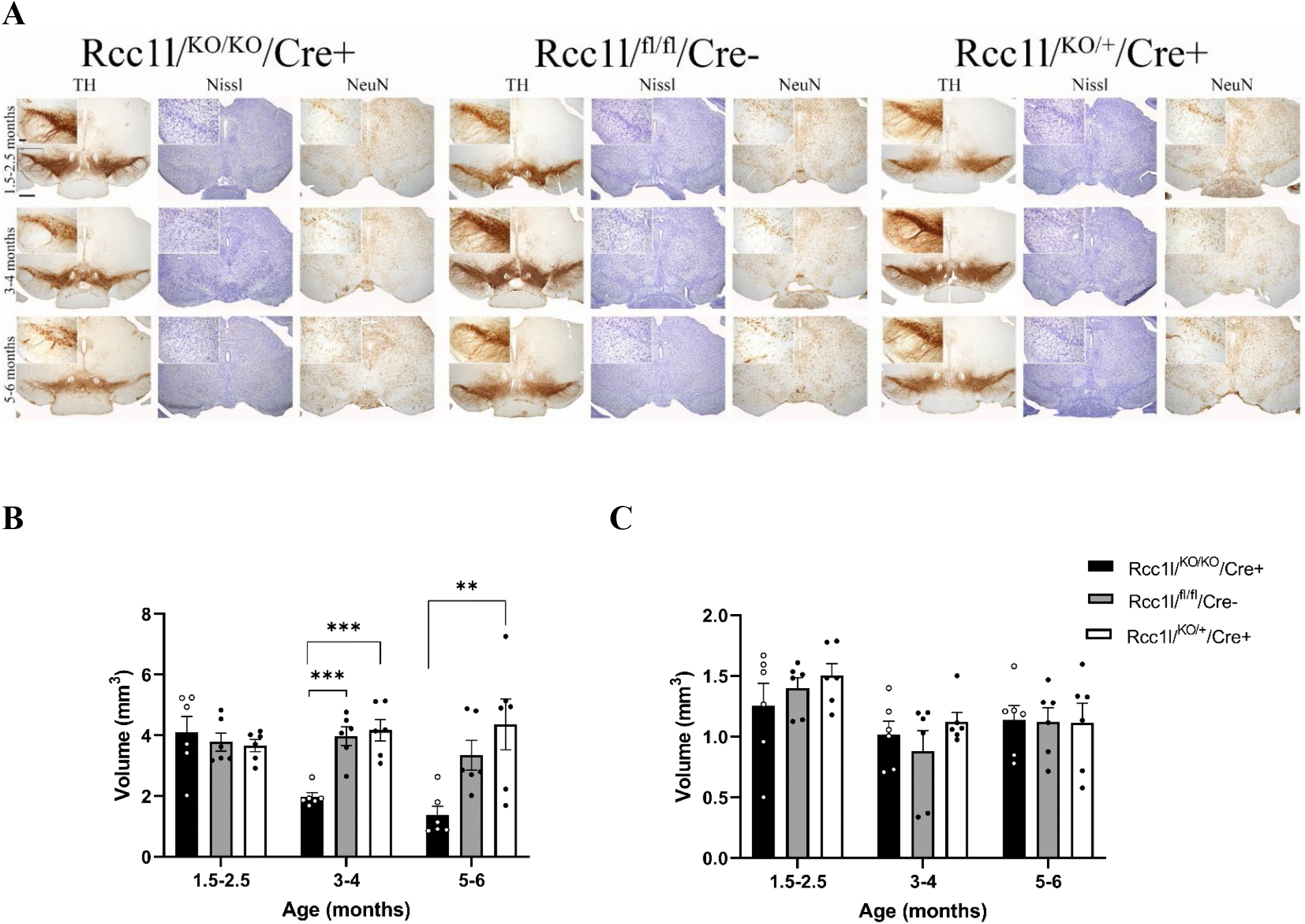
*Rcc1l*^*KO*/*KO*^/*Cre*^+^ mice display an age-dependent reduction of TH immunostaining that is associated with DA neuron loss in the SNc. (A) TH immunostaining reveals a significant loss of evidence of dopaminergic activity in the SNc of *Rcc1l*^*KO*/*KO*^/*Cre*^+^ mice when compared to heterozygotes and wild type littermate controls, beginning at 3-4 months of age. Nissl and NeuN staining demonstrate that loss of TH immunostaining is due to loss of SNc neurons in KO mice, rather than persistence of neurons that have lost the DA phenotype. Top left panels represent a high magnification of the boxed area. Scale bars: 4x, 500 μm and 20x, 100 μm. (B and C) Quantification of the volume of distribution of TH immunoreactivity in the SNc (B), and VTA (C) in the three experimental groups as a function of time (N=3 per experimental group; **, p<0.01, ***, p<0.001).

To determine that a decrease in TH immunoreactivity was not due to loss of dopaminergic phenotype, but instead to neuronal loss, tissue sections were analyzed by Nissl staining and immunohistochemistry for NeuN (Fig. 3A and supplemental figures 2A-4A). Nissl staining reveals a progressive loss of SNc neurons that is confirmed by reduced NeuN staining. Quantification of TH-positive immunostaining was analyzed using densitometry analysis to calculate volume of distribution of TH in the SNc and striatum of the experimental animals. *Rcc1l*^*KO*/*KO*^/*Cre*^+^ mice display significantly reduced TH expression in the SNc when compared to *Rcc1l*^*fl*/*fl*^/*Cre*^-^ and *Rcc1l*^*fl*/*fl*^/*Cre*^-^ controls beginning at 3-4 months of age, as shown by reduced volume of TH immunoreactivity (Fig. 3B; *Rcc1l*^*KO*/*KO*^/*Cre*^+^ *vs. Rcc1l*^*fl*/*fl*^/*Cre*^-^ one way ANOVA main effect of treatment, F _(2,15)_ = 18.76, p=0.0004; *Rcc1l*^*KO*/*KO*^/*Cre*^+^ *vs. Rcc1l*^*KO*/*+*^/*Cre*^+^, p 0.0002) (Figure 3B). No significant loss of TH immunoreactivity was observed in ventral tegmental area (VTA) DA neurons (Fig. 3C). DA projections to the striatum show dramatic reduction in TH immunoreactivity, already striking at 3-4 months of age and even more so at 5-6 months of age (Fig. 4A, and supplemental figures 3B-5B). We also quantified TH immunoreactivity in the striatum to measure the loss of projections from DA neurons in the SNc. Although no significant differences in TH immunoreactivity were observed at the 1.5-2.5-month time point, at 3-4 months of age, *Rcc1l*^*KO*/*KO*^/*Cre*^+^ mice have significantly reduced TH immunoreactivity relative to *Rcc1l*^*fl*/*fl*^/*Cre*^-^ (F _(2,15)_ = 97.23, p<0.0001), and *Rcc1l*^*KO*/*+*^/*Cre*^+^ (F _(2,15)_ = 97.23, p<0.0001) controls (Fig. 4B). A further significant reduction is evident at the 5–6-month timepoint. (*Rcc1l*^*KO*/*KO*^/*Cre*^+^ vs. *Rcc1l*^*fl*/*fl*^/*Cre*^-^, one way ANOVA main effect of treatment F _(2, 15)_ = 242.5, p<0.0001; *Rcc1l*^*KO*/*KO*^/*Cre*^+^ vs. *Rcc1l*^*KO*/*+*^/*Cre*^+^ p<0.0001). Quantification of NeuN immunostaining did not reveal any significant differences between experimental groups at the 1.5-2.5-month time point, at 3-4 months of age, but a significant difference between *Rcc1l*^*KO*/*KO*^/*Cre*^+^ and *Rcc1l*^*fl*/*fl*^/*Cre*^-^ at 5-6 months of age (one way ANOVA main effect of treatment F _(2, 15)_ = 10.44, p= .0014).

**Figure 4.**
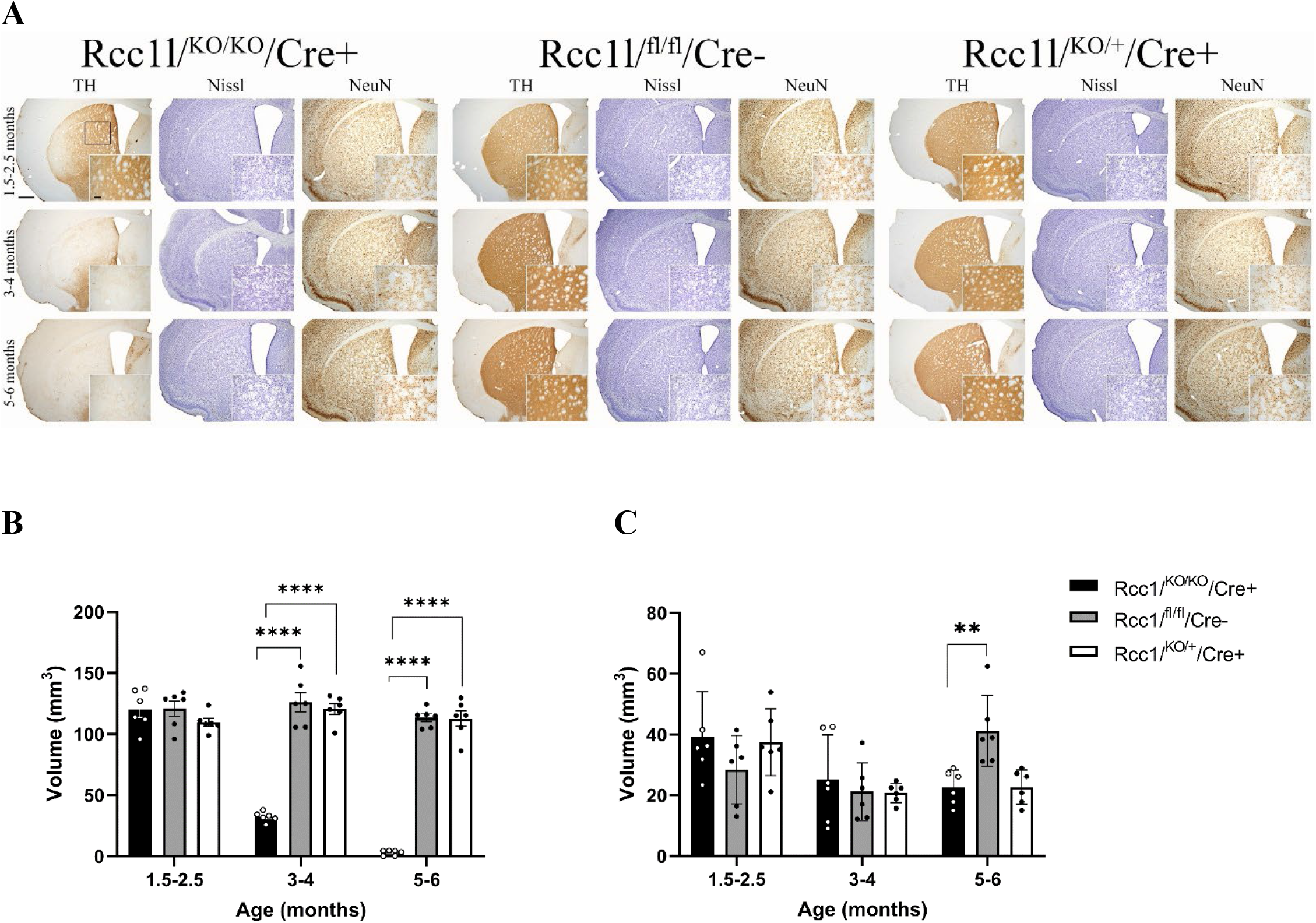
*Rcc1l*^*KO*/*KO*^/*Cre*^+^ mice display an age-dependent reduction of TH positive projections in the striatum. (A) TH immunostaining reveals a significant and dramatic loss of dopaminergic function in the striatum of *Rcc1l*^*KO*/*KO*^/*Cre*^+^ mice when compared to heterozygote and wild type littermate controls beginning at 3-4 months of age. Nissl and NeuN staining demonstrate there are no anatomical differences in the striatum of the different experimental groups. Right bottom panels represent a high magnification of the boxed striatal area. Scale bars: 4x, 500 μm and 20x, 100 μm. (B) Quantification of the volume of distribution of TH immunoreactivity in the striatum in the three experimental groups as a function of time (N=3 per experimental group; ****, p<0.0001). (C) Volume of distribution of NeuN was observed between *Rcc1l*^*KO*/*KO*^/*Cre*^+^ and *Rcc1l*^*fl*/*fl*^/*Cre*^-^ at 5-6 months of age (**, p= .0014).

The dopaminergic VTA projections to the olfactory tubercle and nucleus accumbens seem preserved in *Rcc1l*^*KO*/*KO*^/*Cre*^+^ (supplemental figures 2-4).

### *Rcc1l* mutants display age-dependent mitochondrial fragmentation

We crossed our *Rcc1l*^*KO*/*+*^/*Cre*^+^ mice with the mitochondrially targeted fluorescent reporter mice *Thy1-mitoDendra TM57* (mito-dendra 2)(Magrane et al., 2014) to characterize mitochondria structure in dopaminergic neurons, at the different time points. At 1.5-2.5 months of age, the mitochondria of both *Rcc1l*^*wt/wt*^/*Cre*^+^ and *Rcc1l*^*KO*/*+*^/*Cre*^+^ controls show a diffuse network pattern of distribution in the soma and perinuclear area of DA neurons in the SNc (Fig. 5A) At this time point, some neurons in *Rcc1l*^*KO*/*KO*^/*Cre*^+^ mice have morphologies similar to those in *Rcc1l*^*wt/wt*^/*Cre*^+^ and *Rcc1l*^*KO*/*+*^/*Cre*^+^ controls, but there are already some visibly enlarged spherical mitochondria concentrated around cell nuclei. *Rcc1l*^*KO*/*KO*^/*Cre*^+^ mitochondrial abnormalities are more pronounced at 3-4 and 6 months, showing mostly globular perinuclear aggregates in the few surviving SNc neurons (Fig. 5B).

**Figure 5.**
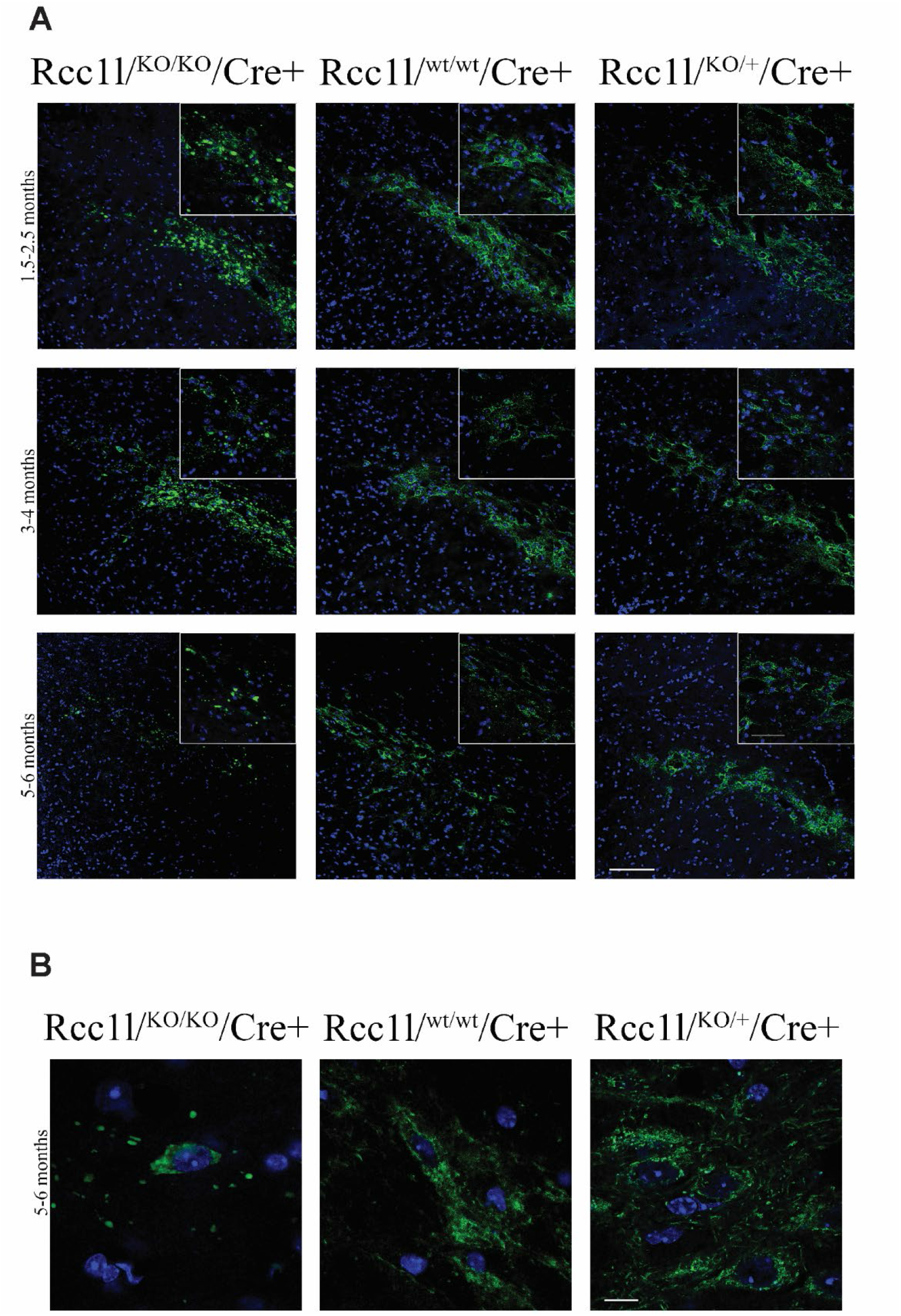
Abnormal mitochondria morphology in the SNc of *Rcc1l*^*KO*/*KO*^/*Cre*^+^ mice. (A) Mitodendra labeling reveals depletion of mitochondria as early as 1.5-2.5 months of age. Enlarged and fragmented mitochondria are visible in the high magnification inset. (B) Perinuclear accumulation of mitodendra labeled mitochondria are apparent in *Rcc1l*^*KO*/*KO*^/*Cre*^+^ mice at 5-6 months of age. Representative images (from N=3 per timepoint/per genotype) were taken from left hemispheres to represent the same area shown in figure 3. Scale bars: 20x, 100 μm; 40x 50 μm; 100x, 50 μm.

## DISCUSSION

This study reveals that loss of protein RCC1L in DA neurons in mice causes onset of a PD-like phenotype that gradually increases in severity. The phenotype includes progressive motor impairments, beginning at 3-4 months of age, associated with significant age-dependent DA neuronal loss in the SNc, and severe loss of DA nerve terminals in the striatum. *Rcc1l*^*KO*/*KO*^/*Cre*^+^ mice exhibit kyphosis and a growth deficit when compared to groups of both heterozygous and wild type littermate controls and die by 6 months of age, even with hydrating gel packs and food provided on the cage floor to compensate for reduced ability to obtain food, due to deficits in locomotion and rearing behavior.

Loss of TH immunoreactivity in the brains of *Rcc1l*^*KO*/*KO*^/*Cre*^+^ mice was accompanied by an apparent a significant decrease in the neuron-specific protein NeuN, and neuronal structure Nissl staining, revealing that DA loss in the SNc of such mice is due to neuronal death, as opposed to loss of dopaminergic phenotype but survival of such neurons (Fig.3 and supplemental figures 2-4). A concomitant progressive and profound loss of projections to the striatum is also observed by TH immunostaining (Fig. 4 and supplemental figures 2-4). Interestingly, we did not observe an intermediate phenotype in heterozygous *Rcc1l*^*KO*/*+*^/*Cre*^+^ mice, suggesting that a single copy of *Rcc1l* may be sufficient to maintain DA neuronal health.

Mitochondrial fusion and fission are required for the establishment of networks, or destruction of damaged mitochondria, respectively. Mitochondrial fusion dysfunction has been associated with PD with the finding that mitochondrial proteins MFN1 and MFN2 are ubiquitinated by PINK/parkin activation (Gegg et al., 2010; Rakovic et al., 2011), and that DA neuron-specific *Mfn2*-null, but not DA neuron-specific *Mfn1*-null, mice display severe loss of DA terminals in the striatum and motor deficits at 5 weeks of age (Lee et al., 2012; Pham et al., 2012). Dopaminergic neurons lacking MFN2 exhibit fragmented/enlarged mitochondria. Drosophila cardiomyocytes lacking Parkin also show enlarged mitochondria (Chen and Dorn, 2013). Mitochondria morphological abnormalities have also been observed in mitochondrial fusion gene OPA1 mutants, where small spherical mitochondria are reported (Griparic et al., 2004). Similar to these findings, we show that mitochondria from dopaminergic neurons lacking RCC1L exhibit morphological abnormalities (Fig. 5). At early time points, fragmented and swollen, spherical mitochondria were found in the SNc of *Rcc1l*^*KO*/*KO*^/*Cre*^+^ mice, similar to the swollen, spherical mitochondria previously found in the DA neurons of DA neuron-specific *Mfn2*-null mice (Lee et al., 2012; Pham et al., 2012), and in contrast to the diffuse weblike pattern of mitochondria observed in wild type and heterozygous controls. The swollen mitochondria found in the SNc of *Rcc1l*^*KO*/*KO*^/*Cre*^+^ mice appear condensed in the perinuclear area, the region in which mitophagy is thought to occur, with targeting of defective mitochondria to this region via mechanisms thought to involve PINK1 and Parkin (Vives-Bauza et al., 2010).

The Lee et al. study (Lee et al., 2012) showed SNc neurons to be viable although their mitochondrial function was deficient, and the DA neuron-specific *Mfn2*-null animals died at 7 weeks of age. In contrast, in the Pham et al. study (Pham et al., 2012), DA neuron-specific *Mfn2*-null animals lived approximately 6 months, or even longer if food was provided in the bottom of the cage, and they showed progressive retrograde loss of striatal terminals and SNc neuronal death. Phenotypic differences between the two studies could be due to the different DAT-*Cre* mice used by Lee et al. (Ekstrand et al., 2007) and Pham et al. (Backman et al., 2006). In support of this possibility, the DA neuron-null *Rcc1l* mice described here employ the same Dat-Cre driver used in the Pham et al study and, like the mice in the latter study, die at approximately 6 months. In fact, the overall similarities between the Pham et al DA neuron-specific *Mfn2*-null phenotype and the DA neuron-specific *Rcc1l*-null phenotype described here are consistent with the possibility of shared *Mfn2, Rcc1l* functions in mitochondrial fusion. It should be noted however, that various features of the DA neuron-null *Mfn2* phenotype in the Pham et al study develop more rapidly than in the *Rcc1l* phenotype, perhaps related to loss of functions that *Mfn2* possesses in addition to roles in mitochondrial fusion. Such functions include roles in axonal transport, and the anchoring of mitochondria to the endoplasmic reticulum, (de Brito and Scorrano, 2008; Misko et al., 2010). Future studies will need to address whether RCC1L is also involved in mitochondrial transport and anchoring. Importantly, as RCC1L has been reported to play roles in intramitochondrial protein translation (Arroyo et al., 2016; Kramer et al., 2023; Reyes et al., 2020), the extent to which deficits in intramitochondrial protein translation may also contribute to the DA neuron-null *Rcc1l* phenotype is also a topic to be addressed by future studies.

The VTA is known to project DA outputs to the nucleus accumbens (NAc) which plays a role in reward processing (Lammel et al., 2014). Surprisingly, the dopaminergic neurons in the VTA and its projections to the VTA as well as the olfactory tubercle (Cleland and Linster, 2019) appear to be preserved in *Rcc1l*^*KO*/*KO*^/*Cre*^+^ mice (Fig.3 and supplemental Figs. 2-4). Pham et al. observed similar TH expression in the VTA of DA neuron-specific *Mfn2*-null mice (Pham et al., 2012). One possible explanation for the preservation of VTA DA neurons is that the DAT-*Cre* driver mouse line employed might display reduced penetrance for targeting DA neurons in the VTA, as has been suggested for some DAT-*Cre* mouse lines (Stuber et al., 2015). This is unlikely in the present study however, as the DAT-*Cre* line used in our study shows robust Cre activity in the VTA (Backman et al., 2006). Another possibility is that VTA DA neurons are more resilient to mitochondrial dysfunction (Pacelli et al., 2015; Ricke et al., 2020; Zampese et al., 2022). The selective vulnerability of SNc DA neurons to neurodegeneration has been a major conundrum in understanding the pathology of PD, since many of the PD causative candidate genes are not expressed solely in SNc neurons. Recent studies point to differences in arborization, with a high number of axonal terminals in SNc relative to VTA, resulting in increased energy demand in the former (Pacelli et al., 2015), and the presence of L-type Ca^2+^ dependent mitochondrial oxidative stress in SNc and not in VTA (Chan et al., 2007; Du et al., 2021; Zampese et al., 2022). Interestingly, as the L-type Ca^2+^ mechanism in SNc DA neurons appears to work via effects on oxidative phosphorylation (Arroyo et al., 2016), and as RCC1-L has been shown to be important to oxidative phosphorylation (Reyes et al., 2020), a mechanism is suggested in which RCC1-L ablation has a particularly deleterious effect on SNc DA neurons. It will be of interest to determine the extent to which SNc and striatal DA loss and PD-like phenotype in *Rcc1l*^*KO*/*KO*^/*Cre*^+^ mice are due to deficits in mitochondrial fusion vs deficits in oxidative phosphorylation, although such studies are complicated by the profound effects that deficits in mitochondrial fusion/fission or in oxidative phosphorylation has on the other process.

Autosomal recessive models null for PINK1(Gispert et al., 2009), Parkin (Perez and Palmiter, 2005), or DJ-1(Goldberg et al., 2005), despite exhibiting some mitochondrial functional abnormalities and subtle deficits in dopaminergic transmission, fail to show nigro-striatal pathology (Dawson et al., 2010; Duty and Jenner, 2011). In fact, triple knockout of PINK1, DJ-1 and parkin together fails to cause loss of DA neurons (Kitada et al., 2009). On the other hand, when PINK/parkin are knocked out in the context of the fission gene knockout Drp1, increased mortality and dramatic loss of DA projections to the striatum is observed (Li et al., 2021). The authors suggest that PINK1 and parkin are not necessary for mitochondria turnover and that only when mitochondria are damaged are these genes important to compensate for an increased need in mitophagy and mitochondrial repair. Indeed, PINK1 KO only showed mitochondrial fission defects under conditions of proteosomal stress (Gispert et al., 2009). In contrast to the PINK1, parkin and DJ-1 models, the *RCCL1* mouse model presented here may be advantageous for certain experimental approaches to studying PD phenotypic traits as it shows pronounced and progressive nigro-striatal pathology even in the absence of further genetic insult.

In conclusion, our study has demonstrated the protein RCCL1 to be vital to the *in vivo* survivability of DA neurons of the nigrostriatal circuit, and that its loss in mice produces a phenotype that includes bradykinesia, postural defects and progressive loss of DA neurons and projections to the ST, which are characteristics of PD progression. Further characterization of this mouse model will determine whether it represents a new model for *in vivo* study of PD, and the putative role of the human *RCC1L* gene as a risk factor that might increase PD occurrence and severity in humans.

## MATERIALS AND METHODS

### Animals

Frozen morula-stage mouse embryos heterozygous for a floxed *Wbscr16*/*Rcc1l* allele (B6Dnk;B6N-Wbscr16^tm1a(EUCOMM)Hmgu^/Ieg) were obtained from the European Mouse Mutant Archive, and were surgically transplanted into pseudopregnant females by standard methodology (Hogan, 1994). *ROSA26::FLPe* mice (Farley et al., 2000), *Slc6a3*^*tm1*.*1(cre)Bkmn*^ mice (Backman et al., 2006) and *Thy1-mitoDendra TM57*(Magrane et al., 2014) were obtained from the Jackson Laboratory. Mice from both sexes were utilized for all experiments described here. Mice were given *ad libitum* access to water and food and maintained on a 12-hour dark-light cycle. All animal procedures were approved by the University of Wisconsin Institutional Animal Care and Use Committee and conducted in accordance with the United States National Institutes of Health *Guide for the Care and Use of Laboratory Animals*. Mice were weighed weekly to monitor their health. When *Rcc1l*^*KO*/*KO*^/*Cre*^+^ mice showed signs of motor impairment (around 4-5 months-old), 2 oz. hydration cups (HydroGel^®^) and food were placed on the cage floor as needed.

For genotyping, ear punches were digested in DirectPCR buffer (Viagen, Cedar Park, TX) with 0.8 mg/ml proteinase K followed by PCR amplification of the resulting DNA. For PCR analysis of excision of the *Rcc1l*^*fl*/*fl*^ locus, primers were: 5’-ACACCACACATCAGTCCATGG-3’ (forward floxed allele-specific), 5’-GGCGCATAACGATACCACG-3’ (forward, excised allele-specific), and 5’-ACTGGAGTCACTTTCCAGGTC-3’ (reverse, common to floxed and excised alleles). For PCR analysis for presence of the *Slc6a3*-*Cre* knocked-in allele, primers were: 5’-TGGCTGTTGGTGTAAAGTGG-3’ (forward, common to both wild type *Slc6a3* and knocked-in *Slc6a3*-*Cre* alleles), 5’-GGACAGGGACATGGTTGACT-3’ (reverse, wild type *Slc6a3*-specific), and 5’-CCAAAAGACGGCAATATGGT-3’ (reverse, *Slc6a3*-*Cre* knocked-in-specific). For PCR analysis of the *Thy1-mitoDendra TM57* knocked-in allele, primers used were: 5’-CTTCCCTCGTGATCTGCAAC-3’ (forward, wild type), 5’-CGCGACACTGTAATTTCATACTG-3’ (reverse, wild type), 5’-GGACATCCCCGACTACTTCA-3’ (forward, *mitoDendra TM57* knocked-in-specific), 5’-GCTCCCACTTCAGGGTCTTC-.3(reverse, *mitoDendra TM57* knocked-in-specific).

### Behavioral Tasks

#### Open Field

All behavioral tasks were carried out during the light phase. Mice were placed individually in a 40.65 × 40.65 × 30.5 cm Plexiglass box filled with 2 cm of corncob bedding and allowed 15 min to explore. Indirect lighting two feet above floor-level was used to encourage normal exploration. Animals were recorded by a ceiling mounted camera using VideoTrack v3.22 (ViewPoint Life Sciences Inc., Montreal, CANADA). Horizontal movement was analyzed using Kinovea v.0.8.15, a 2D motion analysis software (https://www.kinovea.org).

#### Cylinder Test

Forelimb akinesia was evaluated by determining the animal’s ability to support its body weight against the wall of a glass cylinder during exploratory behavior. Mice were placed individually in a transparent glass cylinder (9 cm in diameter and 12.5 cm in height) with corncob bedding covering the bottom. The animals were allowed to explore for 10 minutes while an observer counted the number of weight-bearing touches made with forelimbs to the side of the cylinder.

### Immunohistochemistry

Mice were perfused in 4% paraformaldehyde and brain tissue cryoprotected as previously described (Gerstein et al., 2012). Brains were frozen with dry ice and sectioned in the coronal plane on a sliding microtome at 45 μm thickness into an ethylene glycol-based storage solution and placed at -20°C until ready to use. All processing was carried out at room temperature unless otherwise noted.

Immunostaining against tyrosine hydroxylase (TH) and NeuN was performed as follows. For TH immunostaining **s**ections were washed in 1X TBS followed by antigen retrieval at 80°C in Tris-EDTA pH 9.0 and endogenous peroxidase quenching in TBS with 3% H_2_O_2_ and 10% Methanol. Sections were blocked in TBS with 0.05% Triton X-100 (TBST; Sigma-Aldrich, Burlington, MA) with 5% normal goat serum, incubated overnight in TBST with 1% bovine serum albumin (BSA; Calbiochem by Millipore Sigma, St. Louis, MO) and primary antiserum (Millipore Sigma MAB318; 1:5,000). Following washes in TBST, sections were incubated in 1:300 biotinylated secondary IgG (Vector, Burlingame, CA) for 1 hour, followed by 1 hour in avidin-biotin complex (Standard Elite kit; Vector). Final visualization was obtained incubating in 0.04% 3,3’ – diaminobenzidine (DAB; from tablets; Sigma) and 0.01% H_2_O_2_ in PB, pH 7.4, mounted and coverslipped with Eukitt (Pittsburgh, PA).

For NeuN immunostaining sections were incubated in PBS with 2% BSA and 0.1% saponin (Sigma-Aldrich, St. Louis, MO). Sections were washed, blocked in buffer with 20% normal serum for 45 minutes, incubated overnight in primary antiserum (1:1000, Millipore, MAB377) in 1% normal serum. After washes in PBS+BSA+saponin, sections were incubated in 1:300 biotinylated secondary IgG for 3 hours, followed by 1 hour in avidin-biotin complex. Final visualization was with DAB, mounted and coverslipped with Eukitt.

Additional sections were stained with Nissl as previously described (Hullinger et al., 2013; Osting et al., 2014).

#### Optical Density Analysis

Three animals per experimental group were used for densitometry analysis as biological replicates. Each hemisphere was treated as a technical replicate for statistical analysis resulting in an n=6. A representative series of 45μm sections spanning a total of 5.13mm from striatum through the substantia nigra were used for densitometry measurements. Light microscopic images of matching coronal sections were obtained with a digital camera (Q Imaging Retiga 2000R; Nikon Instruments, Melville, NY) on a Nikon E600W Eclipse epifluorescent microscope with X4-100 planopochromatic objectives with identical light exposure conditions. The optical density of striatum, substantia nigra and VTA was calculated using ImageJ software version 1.53e (NIH ImageJ, Bethesda, MD) by a single, blinded investigator. Each image was converted to an 8-bit grayscale image. ImageJ was calibrated using a step tablet and gray scale values were converted to OD units using the Rodbard function. To minimize the effect of staining differences, background measurements were taken to determine a consistent threshold value and regions of interest (ROIs) were outlined to encapsulate entire brain regions for analysis. ROIs were delineated in a consistent fashion through sections and samples after careful consultation with The Mouse Brain in Stereotaxic Coordinates, (Second Edition. Paxinos and Franklin). OD was measured using the predetermined threshold value defined as the value at which pixels within the sample’s background area remained unlabeled. The size of the area above threshold (AAT) in mm^2^ was determined by the summation of the pixels of ROI staining above the predetermined threshold for all sections measured. Volume above threshold (mm^3^) was calculated by multiplying the AAT value by the number of 45μm sections covering the measured brain region.

### Immunofluorescence

Fixed tissue sections from mice expressing mitoDendra were mounted on subbed slides and coverslipped with ProLong Gold antifade reagent with DAPI. Confocal images were acquired using a Leica SP8 Confocal WLL STED microscope. Fluorescent protein Dendra was excited with a super-continuum white-light laser at 480 nm and emission was detected at 510 nm. Images were obtained with the 20 and 40 X objectives. Images were prepared in Adobe Photoshop. Adjustments on scale, gamma, contrast and hue and subsequent sharpening with the unsharp mask algorithm were applied to the entire image. Montages were assembled with no subsequent retouching. Confocal images were acquired at the focal plane with maximal number of Dendra-labeled puncta from at least three sections per animal and from at least three animals per group.

### Statistical Analyses

Experimenters were blind to the genotype of the animals while performing behavioral testing. Animals from both sexes were used for analysis. Preliminary statistical analysis indicated there were no sex differences in behavioral performance, so sexes were pooled for the final analysis.

All statistical analyses were performed using Prism 9 (Graphpad Software Inc., La Jolla CA) and expressed as mean ± standard error of the mean (SEM). Significance was set at p < 0.05. To analyze behavior and optical density data, one-way ANOVA with Tukey posthoc was used to determine significance between *Rcc1l*^*KO*/*KO*^/*Cre*^+^, *Rcc1l*^*fl*/*fl*^/*Cre*^-^ and *Rcc1l*^*KO*/*+*^/*Cre*^+^ groups at each of the different time points. Mixed-effects analysis was used to analyze weight data, as effect of time and genotype.

## ACKNOWLEDGMENTS

Research reported in this preprint was supported by NICHD of the National Institutes of Health under award number NIH R01 HD079481 to DG, a UW Graduate School Fall Competition grant to DG and R&D Funding from the UW Department of Neurology to CB. We would like to thank Alexander Gowing for excellent technical assistance.

## Notes

**CONFLICT OF INTEREST** The authors have no conflict of interest to report.

### Competing Interest Statement

The authors have declared no competing interest.

### Summary of Updates

Funding support was included. Quantification of NeuN in striatum was added.

