## Supplemental Figures for "Ablation of Mitochondrial RCC1-L Induces Nigral Dopaminergic Neurodegeneration and Parkinsonian-like Motor Symptoms"

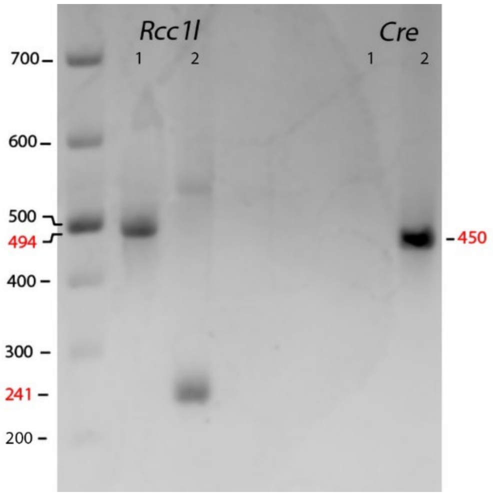


**FIG. S1** Efficient excision of the floxed *Rcc1l* Locus. PCR analysis was performed of ear punch-derived DNA from mice bred to contain either just the floxed *Rcc1l*$^{fl/fl}$ locus (lanes 1) or both the floxed *Rcc1l*$^{fl/fl}$ locus and the Slc6a3-*Cre* knocked-in allele (lanes 2). *Rcc1l*-specific PCR produced 494 bp bands from unexcised *Rcc1l*$^{fl}$ loci and 241 bp bands from excised *Rcc1l*$^{KO}$ loci, respectively, in *Rcc1l*-labeled lanes. Slc6a3-*Cre*-specific PCR produced 450 bp bands from mice with Slc6a3-*Cre* knocked-in allele loci, in *Cre*-labeled lanes.

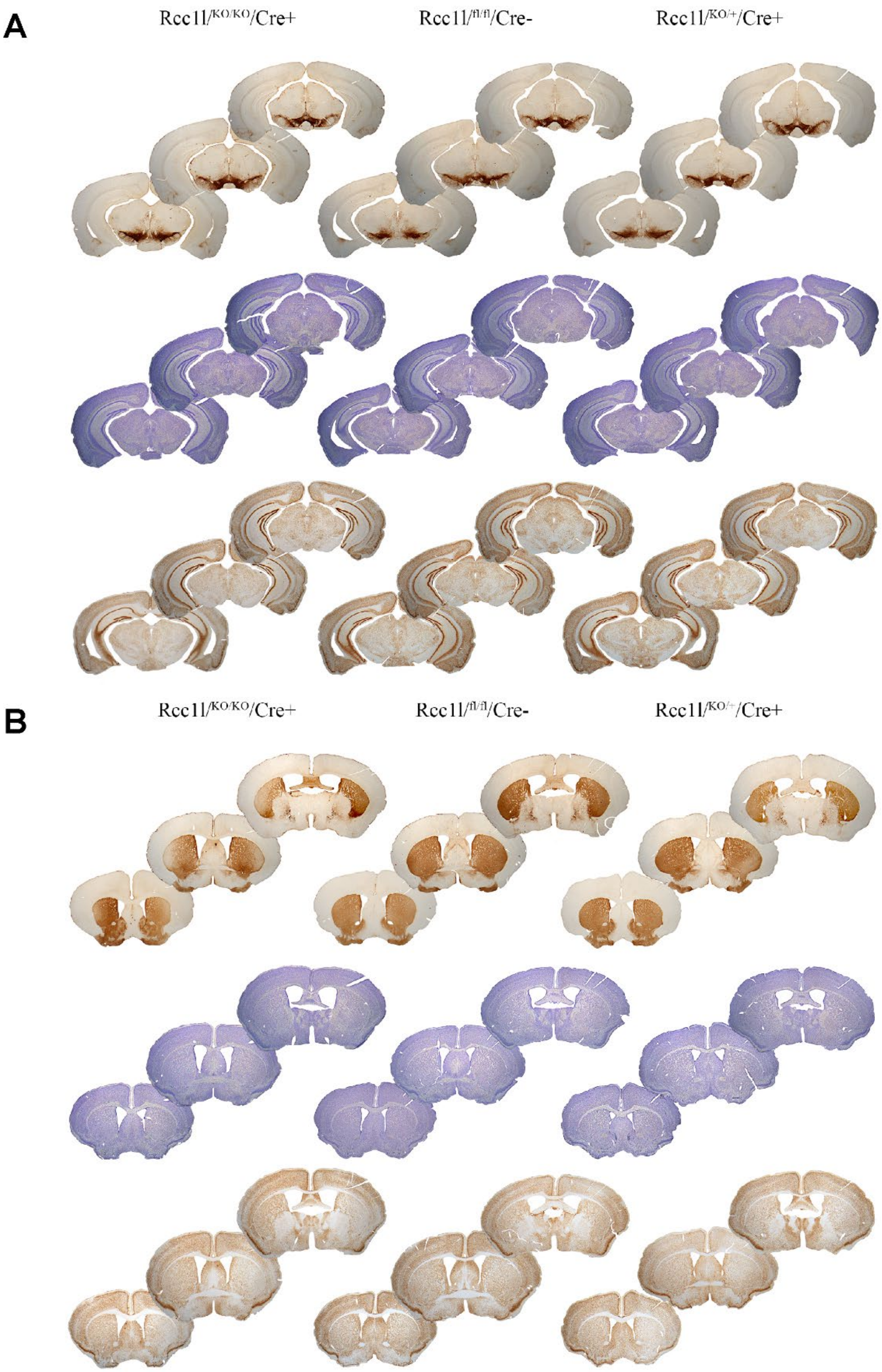


**FIG. S2** Rostrocaudal sections showing extent of TH immunoreactivity (top), Nissl staining (middle) and NeuN (bottom) from the different experimental groups at 1.5-2 months of age (A) Substantia nigra and VTA staining (B) Striatal staining.

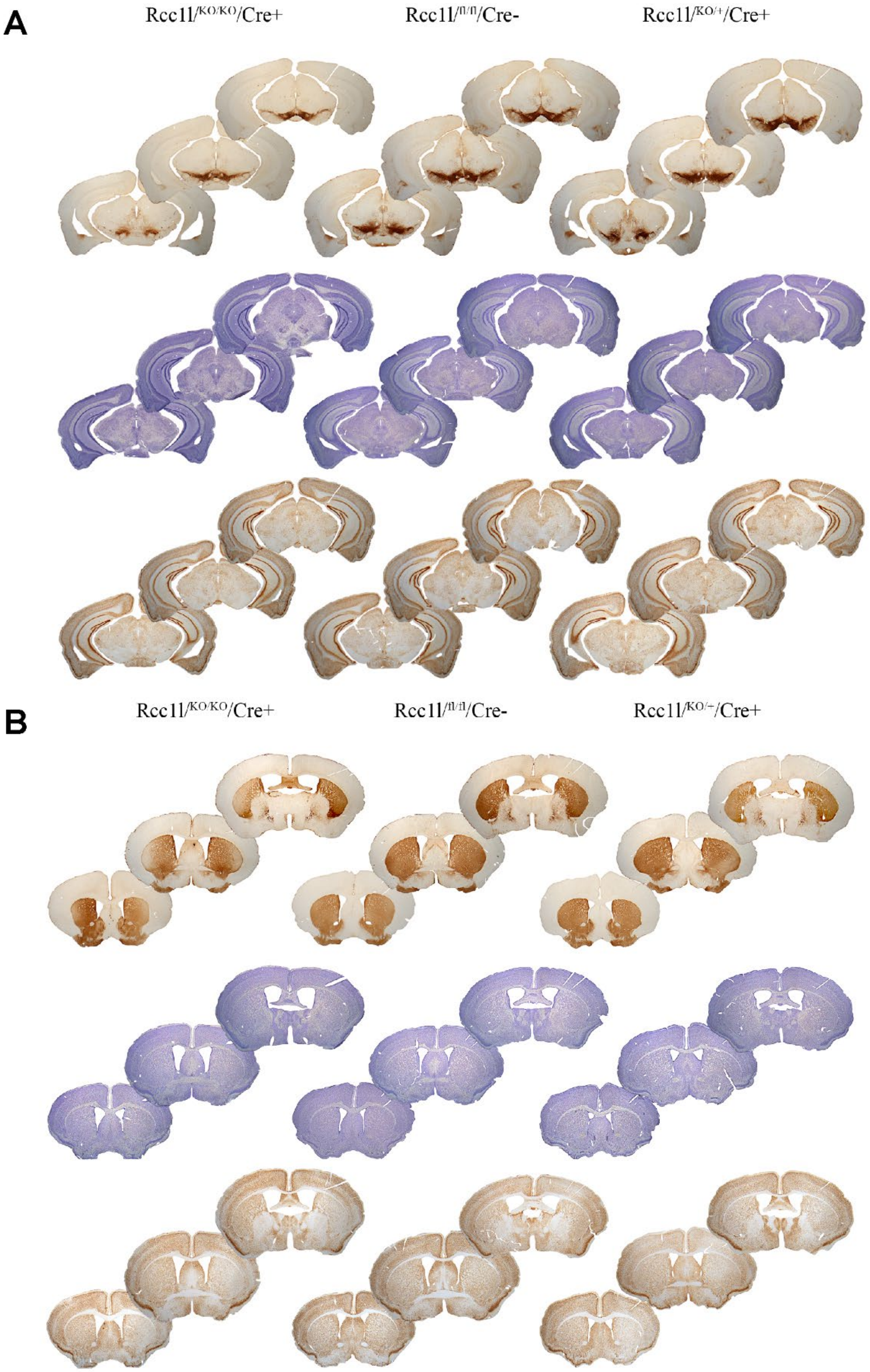


**FIG. S3** Rostrocaudal sections showing extent of TH immunoreactivity (top), Nissl staining (middle) and NeuN (bottom) from the different experimental groups at 3.5-4 months of age
(A) Substantia nigra and VTA staining
(B) Striatal staining.

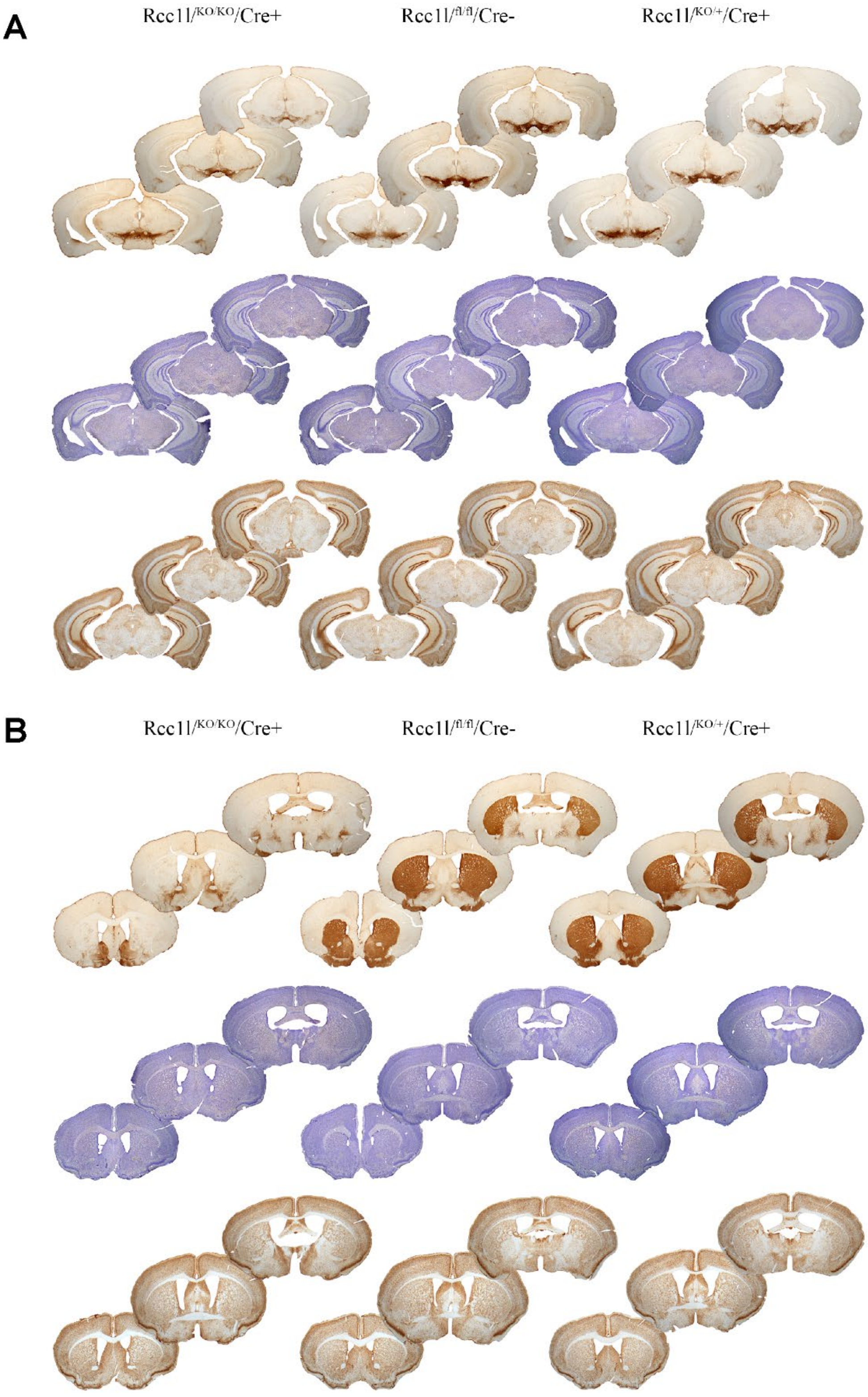


**FIG. S4** Rostrocaudal sections showing extent of TH immunoreactivity (top), Nissl staining (middle) and NeuN (bottom) from the different experimental groups at 6 months of age
(A) Substantia nigra and VTA staining
(B) Striatal staining.
